# Quantitative model of aging-related muscle degeneration: a Drosophila study

**DOI:** 10.1101/2023.02.19.529145

**Authors:** Maria Chechenova, Hannah Stratton, Kaveh Kiani, Erik Gerberich, Alesia Alekseyenko, Natasya Tamba, SooBin An, Lizzet Castillo, Emily Czajkowski, Christina Talley, Anton Bryantsev

## Abstract

Changes in the composition and functionality of somatic muscles is a universal hallmark of aging that is displayed by a wide range of species. In humans, complications arising from muscle decline due to sarcopenia aggravate morbidity and mortality rates. The genetics of aging-related deterioration of muscle tissue is not well understood, which prompted us to characterize aging-related muscle degeneration in *Drosophila melanogaster* (fruit fly), a leading model organism in experimental genetics. Adult flies demonstrate spontaneous degeneration of muscle fibers in all types of somatic muscles, which correlates with functional, chronological, and populational aging. Morphological data imply that individual muscle fibers die by necrosis. Using quantitative analysis, we demonstrate that muscle degeneration in aging flies has a genetic component. Chronic neuronal overstimulation of muscles promotes fiber degeneration rates, suggesting a role for the nervous system in muscle aging. From the other hand, muscles decoupled from neuronal stimulation retain a basal level of spontaneous degeneration, suggesting the presence of intrinsic factors. Based on our characterization, *Drosophila* can be adopted for systematic screening and validation of genetic factors linked to aging-related muscle loss.

## INTRODUCTION

Virtually all animals demonstrate some degree of somatic muscle tissue loss during aging. Muscle mass or size decline has been reported for a diverse range of organisms, from worms to mammals [1–4]. In older humans, the condition of aggravated loss of muscle mass, accompanied by a profound decrease of muscle power, is known as sarcopenia [5]. World-wide, sarcopenia affects 10% of community-living individuals older than 60 years [6]. Complications arising from sarcopenia negatively impact the quality of senior living, contribute to increased morbidity and mortality, and bear billions of dollars in healthcare costs [7–9].

Muscle mass is reflective of changes in the size and number of individual cellular units, muscle fibers. Atrophic changes affect the size of muscle fibers and in the elderly these changes can be reversed by physical exercise [10]. In contrast, dystrophic, degenerative changes are not likely to be reversed because of the physical elimination of muscle fibers. A significant loss of fibers has been detected by whole-muscle quantification of the leg muscles in old humans and mice [3, 11, 12]. Although mammalian muscle fibers have a strong regenerative potential for acute injury [13], this capacity may decline upon chronic injury or with advanced age [14, 15]. Therefore, the cumulative effect of degenerative changes in muscle fibers is a considerable factor for the decline of muscle mass in older humans [16, 17].

Multiple studies assessed muscle size and strength in young and old subjects and confirmed a strong genetic determination of these traits [18]. However, the genetics of sarcopenia remains difficult to dissect, which is highlighted by regular ambivalence in reported results [18]. Currently, most of the studies looking for genetic variants associated with sarcopenia operate with rather a limited set of pre-selected candidate genes that have been chosen based on their involvement in general muscle development and maintenance [19–21]. Ideally, these approaches should be complemented by studies that use an unbiased approach to identify novel genetic factors that contribute to muscle aging.

The fruit fly *Drosophila melanogaster* is one of the leading model organisms in genetic research, offering a well-developed research toolbox for genetic screening, complete with rigorous validation approaches [22, 23]. Being amenable to genome-wide, systematic screening, the *Drosophila* model provides significant insights into genes that are involved in muscle development and functionality [24, 25]. Flies also have been informative for identifying genetic modifiers of simulated human degenerative muscle conditions [26, 27]. With the development of the Drosophila genetic research panel (DGRP), consisting of >200 fully sequenced isogenic fly lines, it becomes very possible to run genome-wide association studies to probe complex, multifactorial traits, such as longevity, behavior, etc. [28–30]. Selective genetic drivers can allow gene manipulation in specific muscle groups, while monitoring the outcomes via functional tests [31]. However, the major roadblock preventing full adoption of the *Drosophila* model for muscle aging research is the lack of knowledge about muscle changes that occur in aging flies.

In this study, we provide the initial characterization of aging-related muscle degeneration in *Drosophila*. We demonstrate that spontaneous degeneration of muscle fibers occurs in all types of somatic muscles of adult flies. Using quantitative approaches, we show that this process is strongly correlated with chronological and functional aging of flies. Further, we apply experimental genetics to test how mechanical overloading and functional inactivation of muscles can affect the rate of fiber degeneration during aging. Our findings enable the *Drosophila* model for systematic analysis of genetic factors contributing to aging-related muscle loss.

## MATERIALS AND METHODS

### Fly stocks and husbandry

Flies were cultured on Jazz mix food (Fisher) on a 12-h light/dark cycle. For aging studies, 1-2 day old adults were placed in standard plastic vials (Genesee Scientific), not more than 35 flies/vial, and kept at 29°C; vials were changed twice a week. For aging females, three male mating partners were additionally added per vial. Most fly lines were supplied by the Bloomington Drosophila Stock Center (BDSC); **Table 1** lists all genetic lines used in this study. Genetic crosses were set up at 25°C and newly eclosed adults were transferred to 29°C for aging. A control cross for RNAi knockdown experiments was set between a driver line and *attP2* flies for the best genetic matching (**Table 1**). The mechanical stimulation applied in the experiments involving bang-sensitive flies was described in *Horne et al* as “vortex testing” [32]. In brief, a standard culture vial containing flies was shaken for 10 sec using a lab vortexer set to maximum speed. Such treatment was repeated daily for the entire duration of aging trial (4 weeks).

**Table 1.** The specifics of genetic lines used in this study.

| Short ID | Source | Genetic background | Notes |
| --- | --- | --- | --- |
| *w<sup>1118</sup>* | unknown | w[1118] |  |
| *yw* | unknown | y[*] w[*] |  |
| DGRP-208 | BDSC 25174 | wild-type |  |
| DGRP-301 | BDSC 25175 | wild-type |  |
| DGRP-303 | BDSC 25176 | wild-type |  |
| DGRP-304 | BDSC 25177 | wild-type |  |
| DGRP-307 | BDSC 25179 | wild-type |  |
| DGRP-313 | BDSC 25180 | wild-type |  |
| DGRP-324 | BDSC 25182 | wild-type |  |
| DGRP-357 | BDSC 25184 | wild-type |  |
| DGRP-360 | BDSC 25186 | wild-type |  |
| DGRP-820 | BDSC 25208 | wild-type |  |
| DGRP-852 | BDSC 25209 | wild-type |  |
| DGRP-705 | BDSC 25744 | wild-type |  |
| DGRP-21 | BDSC 28122 | wild-type |  |
| DGRP-41 | BDSC 28126 | wild-type |  |
| DGRP-45 | BDSC 28128 | wild-type |  |
| DGRP-59 | BDSC 28129 | wild-type |  |
| DGRP-75 | BDSC 28132 | wild-type |  |
| DGRP-88 | BDSC 28135 | wild-type |  |
| DGRP-91 | BDSC 28136 | wild-type |  |
| DGRP-93 | BDSC 28137 | wild-type |  |
| DGRP-105 | BDSC 28139 | wild-type |  |
| DGRP-136 | BDSC 28142 | wild-type |  |
| DGRP-142 | BDSC 28144 | wild-type |  |
| DGRP-149 | BDSC 28145 | wild-type |  |
| DGRP-153 | BDSC 28146 | wild-type |  |
| DGRP-158 | BDSC 28147 | wild-type |  |
| DGRP-161 | BDSC 28148 | wild-type |  |
| DGRP-85 | BDSC 28274 | wild-type |  |
| DGRP-57 | BDSC 29652 | wild-type |  |
| 79B6 | [33] | w[1118] P( w[+mC]=Act79B-Gal4) | TDT-specific driver for the Gal4/UAS system, employed in KD<sup>TDT</sup> flies. |
| *elav-Gal4* | BDSC 458 | P([+mW.hs]=GawB)elav[C155] | neuron-specific driver for the Gal4/UAS system, employed in KD<sup>NEU</sup> flies. |
| *Mef2-Gal4* | Gift from Dr. Aaron Johnson, Washington University in St. Louis | w[*];P(w[+m*]=Mef2-Gal4)3 | Muscle-specific driver for the Gal4/UAS system, used in KD<sup>MUS</sup> flies. |
| *Ca-α1D KD* | BDSC 33413 | y[1] sc[*] v[1]; P(y[+t7.7] v[+t1.8]=TRiP.HMS00294)attP2 | induces *Ca-α1D* RNAi knockdown |
| *jus KD* | BDSC 28764 | y[1] v[1]; P(y[+t7.7] v[+t1.8]=TRiP.JF03192)attP2 | induces *jus* RNAi knockdown |
| *attP2* | BDSC 36303 | y[1] v[1]; P(y[+t7.7]=CaryP)attP2 | control for RNAi knockdown crosses |

### Functional tests

For the jump test, flies with clipped wings were tested on a sheet of plain paper. The flies were stimulated to jump by gently touching the abdomen with a paintbrush; the takeoff and landing spots were marked by pencil and then the distance between the marked spots was measured with a ruler. The average distance obtained from three jumping trials was recorded for each fly. For the flight test, flies were released one-by-one in the center of a graded flying chamber and their landing location was used to classify each fly as flying upward (U), horizontally (H), downward (D), or not flying at all (N). The flight index was calculated according to the formula: (6x[U flies]+4x[H flies]+2x[D flies]+0x[N flies])/[all tested flies] [34]. For the climbing test, batches of 10 flies were placed into a graded standard fly vial without food and gently tapped while being recorded with a smartphone camera in slow-motion mode. The number of flies that reached height thresholds (set to 1, 4, and 7 cm) in 10 sec after the tapping was noted.

### Cryosectioning and immunofluorescence

We followed general guidelines of sample preparation, as previously described [35]. Flies were sectioned in the horizontal plane, at 7-10 µm thickness and air-dried on standard microscopy slides (SuperFrost Plus, Fisher) We used the following mouse monoclonal antibodies from Developmental Studies Hybridoma Bank: anti-Dlg (clone 4F3), anti-α integrin (clone DK.1A4), and anti-*β*-integrin (clone CF.6G11). Incubation with primary antibodies (diluted 1:50) was done overnight in a staining solution (Phosphate Buffered Saline (PBS) supplemented with 0.1% Triton X-100 and 1% Bovine Serum Albumin (BSA)), using a humid chamber at room temperature. For anti-integrin staining, the two monoclonal antibodies were combined and applied simultaneously. The secondary antibody was Cy3-labeled goat anti-mouse (115-167-003, Jackson ImmunoResearch), incubated in the staining solution (without BSA) for 1 h at room temperature. For muscle and nuclear counterstaining, we used phalloidin conjugated with iFluor 488 (ab176753, Abcam) and DAPI (Sigma), respectively.

### Abdominal preparations

For abdominal wall muscles, whole flies were fixed in PBS-buffered 4% formaldehyde overnight at 4°C and then dissected. Using 22 G syringe needles (BD305155, VWR), the abdomens were cut off and opened up with a longitudinal incision along the dorsal side. After removal of internal organs, the abdominal walls were stained with phalloidin and DAPI for 1 h at room temperature and then mounted with a coverslip on a standard microscopy slide for imaging.

### Histochemistry

We essentially followed a previously published protocol [36, 37]. In brief, microscopy slides containing fresh, 15-µm cryosections were incubated at room temperature with 200 µL/slide of the staining solution (50 mM Tris pH7.4, 1 mg/mL nitro blue tetrazolium chloride, 5 mM MgCl2, 50 mM sodium succinate, 10 mM sodium azide). Sections from young and old flies were stained in parallel. Staining progress was monitored under a dissecting microscope; reactions were terminated by moving the slides into the fixing solution (4% formaldehyde in PBS) for 1o min.

### Microscopy and image acquisition

AxioImager 2 (Zeiss) equipped with 20X (0.8 NA) objective and color and monochrome CCD cameras (Axiocam HR and Axiocam MR, Zeiss) was used for routine examination of slides. Select samples were imaged with the laser confocal microscope LSM 700 (Zeiss). Image acquisition was done via the Zen software (Zeiss). Image cropping and image intensity adjustments for figures was done via Photoshop (Adobe).

### Quantification of muscle fiber loss

Typically, 15 flies (*i*.*e*. 30 TDTs or subalar muscles) were analyzed per sample. Every fluorescently labeled muscle was imaged 2 or 3 times from tissue sections obtained at various depths within the thorax. Two human operators independently analyzed the images to count intact (live) fibers and to estimate degeneration scores. Special care was taken to recognize and exclude from the analysis artifacts caused by mechanical damage, such as tissue tears. If two images of the same muscle were not concordant on the severity of degeneration, the more severe damage was recorded. To calculate the mean of aging-related changes in live TDT fibers 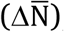, the mean of fiber counts obtained from the young flies 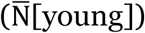 was subtracted from individual fiber counts (N_1_[old]… N_n_[old]) obtained from old TDTs (n) and then the average of individual differences was calculated, according to the formula: 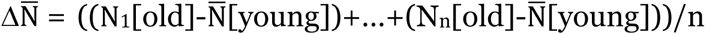.

For the degeneration score system, muscles with all fibers intact received score “0”, muscles with a single degenerate or dead fiber received score “1”, muscles with multiple degenerate fibers (typically 2-4) received score “2”, and those muscles that were either no longer holding together as a whole or where the damage affected >50% of the cross- sectioned area received score “3”. The distribution of scores within a sample in graphs was depicted by circles which areas represented the ratio of muscles received each penalty score; the sum of all circles makes 100%, *i*.*e*. all muscles analyzed in the sample.

### Statistical analysis and calculations

Box plots were created with the online tool [38]. Statistical significance between control and experimental groups was calculated using the Student’s *t-*test. Differences were deemed statistically non-significant with *p>0*.*05*. Statistically significant differences were denoted by * (for *p<0*.*05*) and ** (for *p<0*.*01*).

## RESULTS

### Functional decline of muscle performance in aging flies

For the initial tests, we chose *w*^*1118*^ as this typical lab stock line has been used as parental line for tens of thousands of transgenic lines (>25,000 listed by BDRC) and often serves as comparative negative control in experiments involving transgenics. We opted to conduct aging at 29°C as it constitutes the upper limit of physiologically relevant temperature zone for *D. melanogaster* [39]. The higher temperature notably reduces the duration of aging trials and increases experimental turnaround.

Populations of *w*^*1118*^ flies reared at 29°C gradually declined and became completely extinct within 6 weeks (**Fig. 1A**). For further analyses we selected three ages (0, 2, and 5 weeks) as checkpoints through *w*^*1118*^ lifespan. Muscle functionality was assessed by three tests that independently probed jump, flight, and leg musculature, respectively. The jumping ability declined precipitously in 5 w.o. flies (**Fig. 1 B, C**). The ability to fly (**Fig. 1D**) and climb (**Fig. 1E**) was declining more gradually but also reached the absolute minimum in 5 w.o. flies.

**Figure 1:**
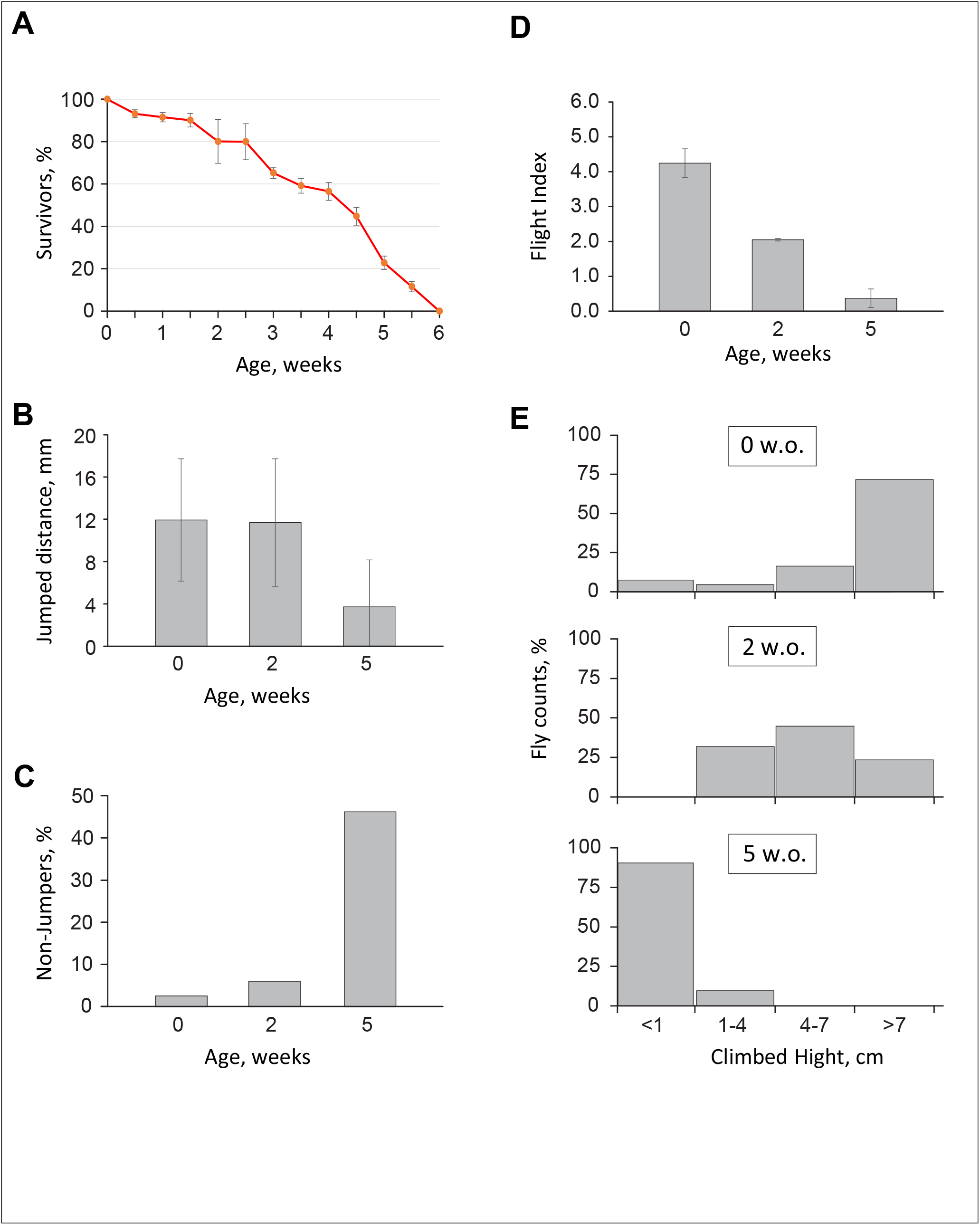
Functional decline of somatic muscles in aging *w*^*1118*^ flies. **A**. Populational changes of the *w*^*1118*^ flies assessed over the lifetime. The data were collected from 3 independent batches of 100 *w*^*1118*^ flies reared at 29°C. **B**. The average distances jumped by a *w1118* fly at select ages. The data were obtained from 35-50 *w*^*1118*^ flies per age group. Error bars represent standard deviation. **C**. The percentage of flies that cannot jump. The data were obtained from 35-50 *w*^*1118*^ flies per age group. **D**. The flight ability of the *w*^*1118*^ flies at select ages. The flight index [34] was calculated from 60-120 flies per age group. Error bars represent standard deviation. **E**. The climbing performance at select ages. The data represent distribution of climbed heights achieved by 50-60 *w*^*1118*^ flies in a 10 sec trial.

We concluded that, based on the functional criteria, *w*^*1118*^ flies reared at 29°C become old at the age of 5 weeks. We further hypothesized that old flies may have accumulated changes in their muscles that prevented them from optimal performance in the functional tests.

## Instances of spontaneous muscle degeneration in aging flies

To detect degenerative changes in somatic muscles, we relied on universal vital and structural markers. Succinyl dehydrogenase (SDH) activity represents intact, functional mitochondria and serves as a classical marker of cell viability. We studied SDH activity in unfixed cryosections of flies, using histochemistry [36, 37]. Indirect Flight Muscles (IFMs) as well as many smaller thoracic muscles (*e*.*g*., subalar muscles) showed strong SDH activity, while degenerate or dying muscle fibers completely lost it (**Fig. 2A**). Using SDH activity staining, we detected multiple instances of fiber degeneration in IFMs and other oxidative muscle fibers in 5 w.o. flies, however this method was not universal, because some muscles (*e*.*g*., the Tergal Depressor of the Trochanter, TDT, also known as jump muscle) naturally had a very low basal SDH activity levels (**Fig. 2A**).

**Figure 2:**
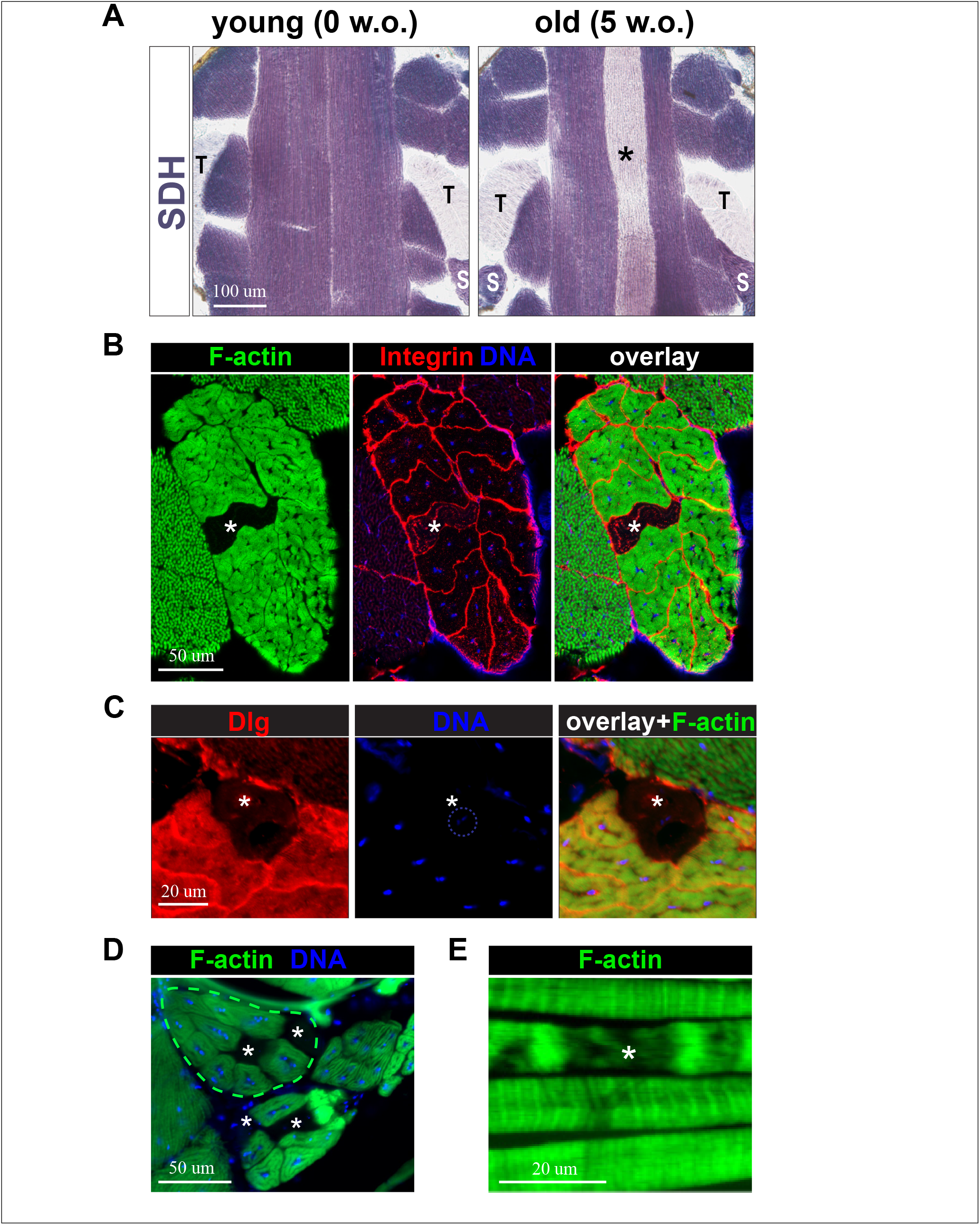
Spontaneous muscle fiber degeneration in aging *w*^*1118*^ flies. **A**. Thoracic muscles stained histochemically for SDH activity. Oxidative IFM and subalar (S) muscles are stained dark, glycolytic TDT muscle (T) remains pale. A degenerate IFM fiber (asterisk) loses the SDH staining. Anterior is on top. **B**. A cross-sectioned TDT muscle from an old fly. Individual TDT fibers are visualized by integrin staining (red); a degenerate fiber (asterisk) loses the presence of F-actin (green) and DNA (blue). **C**. A magnified anterior edge of the TDT muscle from an old fly. A degenerate, dying fiber (asterisk) is missing a fine Dlg-positive T-tubule network (red), F-actin (green) and has a severely diminished DNA content (blue, circled). **D**. The instance of multiple degenerate fibers in the subalar muscle (outlined with a dashed line) as well as other small thoracic muscles. Degenerate muscle fibers that are missing F-actin (green) and DNA (blue) are marked by asterisks. **E**. Segmental degeneration within the lateral abdominal muscles. The asterisk marks a degenerating segment of a muscle that lacks F-actin content.

To assess fiber viability in the TDT muscle, we analyzed the structural component of muscle fibers. The TDT is a composite muscle consisting of multiple individual fibers that contain filamentous actin (F-actin) at high density (**Fig. 2B**). In 5 w.o. *w*^*1118*^ flies, we observed random TDT fibers that lost F-actin (**Fig. 2B**). Upon further examination, such fibers were also found to lose the intracellular network of T-tubules (revealed by anti-Dlg immunostaining) and nuclear DNA (**Fig. 2C**). Since these changes are detrimental to muscle functioning and viability, we concluded that muscle fibers with no F-actin content must be degenerate, dying or already dead. We did not detect cellular and nuclear fragmentation in degenerating fibers.

Individual fibers with a loss of F-actin were observed in all muscle groups of adult flies, including small thoracic muscles and abdominal wall muscles (**Fig. 2 D,E**). The latter also demonstrated that F-actin loss could be local, affecting only a segment within a muscle fiber (**Fig. 2E**). Sometimes, segmental degeneration was also observed on serial sections of the TDT muscle (not shown).

Collectively, morphological analysis revealed that old *w*^*1118*^ flies contain degenerate muscle fibers. Meanwhile, after having examined many samples, we noticed that muscle degeneration is stochastic as not every old fly had degenerate fibers. To establish a link between aging and muscle degeneration in flies, we conducted a quantitative study.

## Quantification of muscle degeneration

The TDT muscle was chosen for fiber loss quantification because of its large size and multi-fiber composition. In this study we used flies of advanced age (5 w.o.) and included another common laboratory strain, *y w*. As previously reported [40], the average fiber content per TDT was line-specific, although fiber counts in individual flies varied slightly (**Fig. 3A**). There was a statistically significant reduction in the counts of live TDT fibers from young and old flies of both genetic lines (**Fig. 3A**).

**Figure 3:**
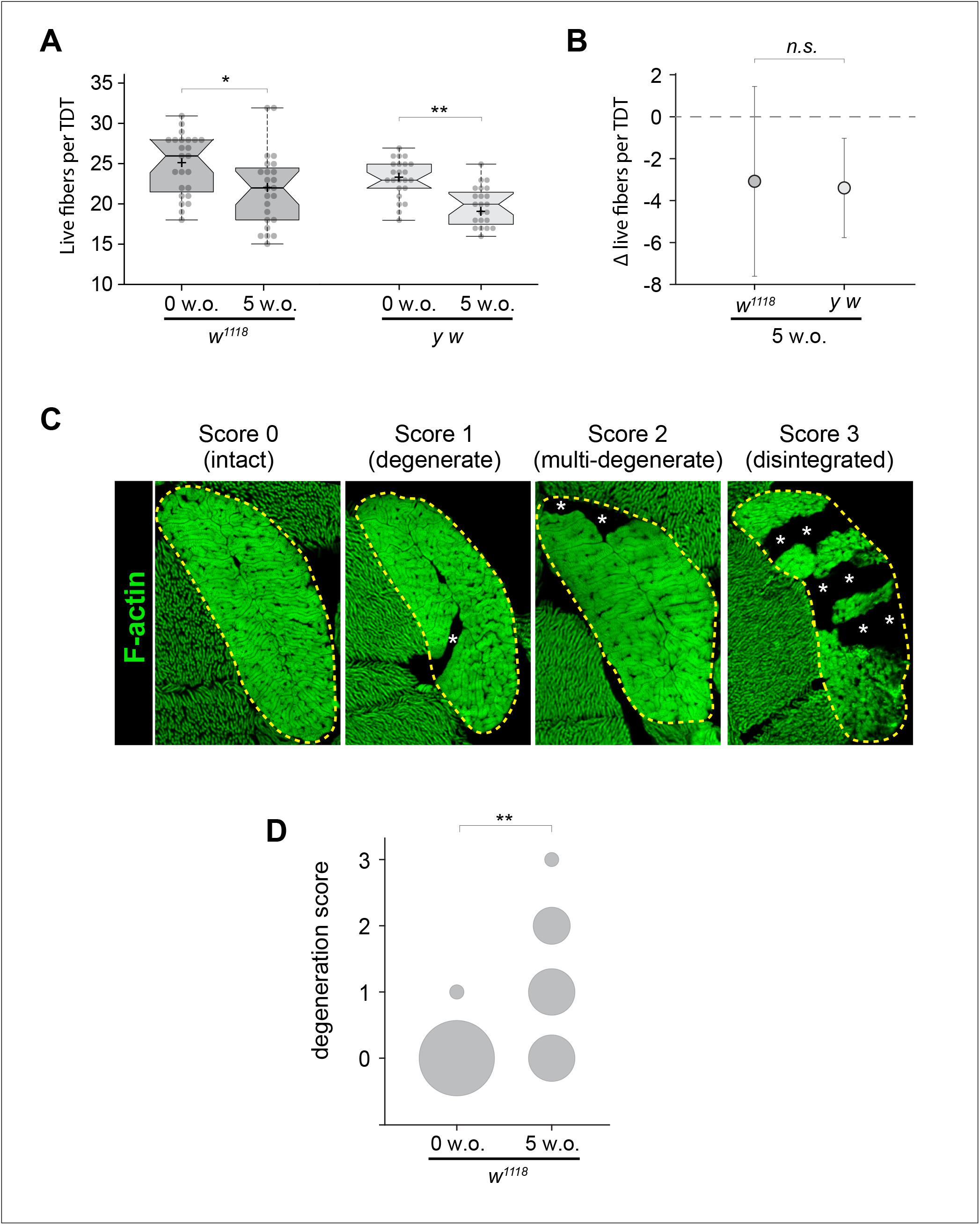
Quantification of spontaneous fiber degeneration in the TDT muscle in aging flies. **A**. Box plots of live fiber counts obtained from the TDTs of *w*^*1118*^ and *y w* flies aged at 29°C for 3 days (0 w.o.) and 5 weeks (5 w.o.). Each box plot is based on data obtained from 20-25 TDT muscles. **B**. The relative changes in live TDT fibers in old *w*^*1118*^ and *y w* flies. The same data input was used as in A. Whiskers show standard deviation, dotted line sets the value of young flies (0 w.o.). **C**. Examples of TDT degeneration severity along with corresponding penalty scores. The TDT muscle is outlined with a dashed line, asterisks indicate places of muscle degeneration. Images represent a single optical section obtained with the laser confocal microscope. **D**. Quantification of the TDT damage extent in young (0w.o) and old (5 w.o.) *w*^*1118*^ flies. The area of each circle represents the fraction of muscles received a particular penalty score. Values of the Student’s *t-*test: * *p<0*.*05*; *** *p<0*.*001*; *n*.*s*.: *p>0*.*05*.

Line-specific differences in the mean TDT fiber counts complicated direct comparison of data obtained from different genetic lines. To normalize data, we expressed the loss of live TDT fibers for each line as a difference between the counts obtained from old flies and young flies. After such normalization, the extent of fiber loss became comparable across different lines (**Fig. 3B**).

From a practical standpoint, analysis of live fibers is taxing because it requires two sets of samples (young and old) and generates high volume of data. To streamline quantification of degeneration, we applied a system of penalty scores reporting the severity levels of fiber degeneration (**Fig. 3C**).

According to penalty-based assessment system, old flies demonstrate significantly elevated damage in their TDTs (**Fig. 3D**). This approach does not require input from young flies, has a faster turnaround, and generates data that is readily compatible across different genetic lines.

Overall, by applying different approaches we demonstrate that the severity of muscle degeneration can be quantified and cross-compared between genetic lines.

## Spontaneous muscle degeneration is aging-related

We quantified the extent of TDT degeneration in aging *w*^*1118*^ flies over their entire lifespan. The counts of live TDT fibers declined steadily, reaching statistically significant differences at advanced ages (**Fig. 4A**). The change in live TDT fibers strongly and inversely correlated with the chronological age of flies (correlation coefficient *r*= −0.93).

**Figure 4:**
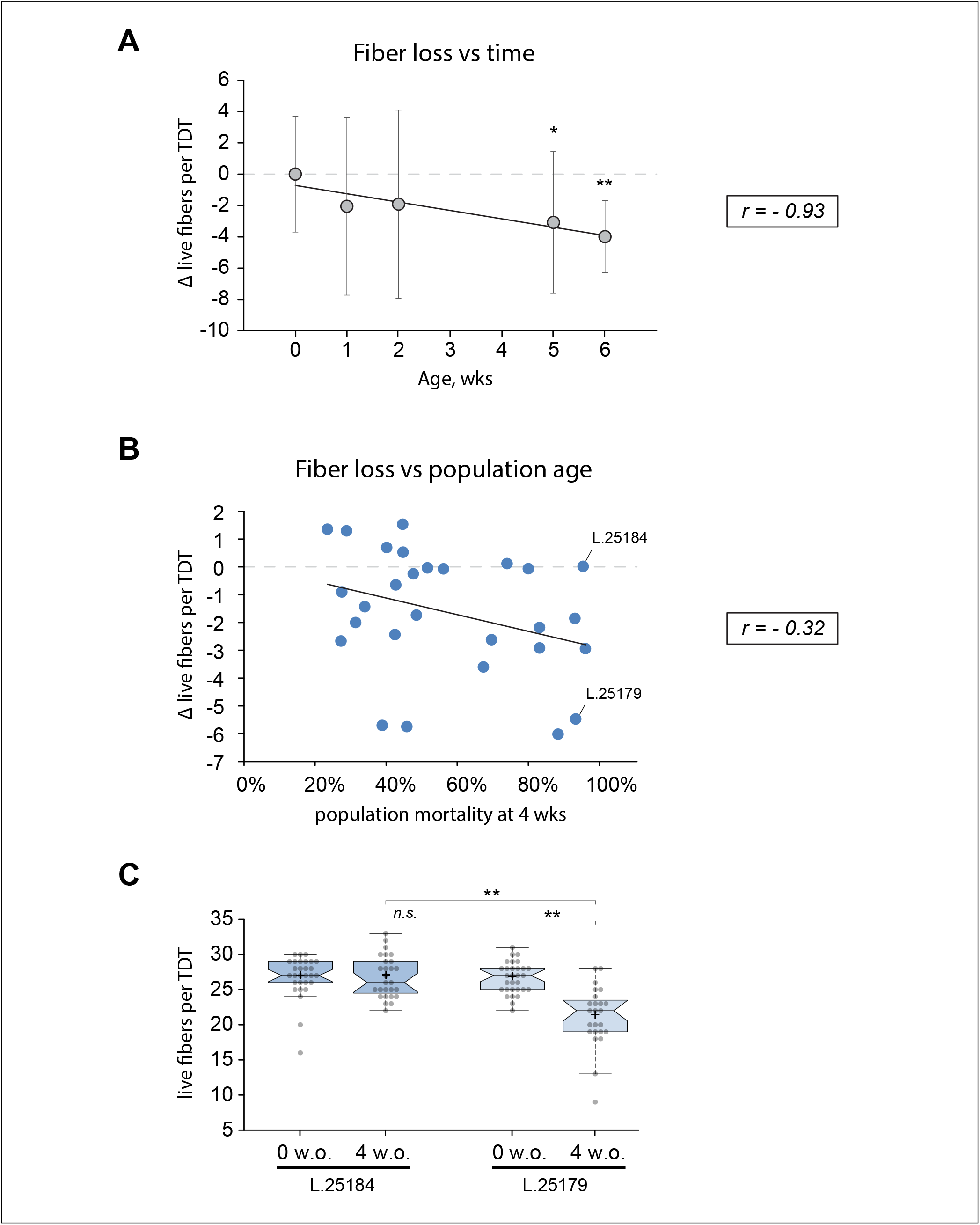
Spontaneous muscle fiber loss in flies is aging-dependent and is influenced by genetic background. **A**. Correlation between chronological age and changes in live TDT fibers counts obtained from *w*^*1118*^ flies reared at 29°C. Each point represents data from 25-30 TDTs (10 TDTs were used at 6 weeks). Whiskers show standard deviation. Dashed line marks the value of young flies (0 w.o.). Solid line depicts linear regression with the correlation coefficient (r) displayed. **B**. Correlation between population age and changes in live TDT fiber counts of 29 inbred wild-type DGRP lines. Population ages are expressed based on mortality percentile of 100 flies aged at 29°C for 4 weeks. L.25184 and L.25179 exemplify lines with similar mortality rates but different TDT degeneration rates. Other labels are as in A. **C**. Box plots of live TDT fiber counts in two exemplary fly lines (labeled in B) at the beginning (0 w.o.) and end (4 w.o.) of the aging trial. Notches represent the 95% confidence interval. Values of the Student’s *t-*test: * *p<0*.*05*; *** *p<0*.*001*; *n*.*s*.: *p>0*.*05*.

The link between TDT fiber losses and aging was further tested with multiple isogenic wild-type lines from the Drosophila Genomic Reference Panel (DGRP, [29]). To accommodate for putative lifespan differences of individual lines, all DGRP lines were aged for a fixed amount of time (4 weeks) and their physiological age was estimated as population mortality at the end of the trial period. Based on the results from 29 tested lines (**Fig. 4B**), the extent of TDT fiber loss remained in a solid inverse correlation with age (*r*=-0.32).

Individually, DGRP lines demonstrated a wide range of degeneration rates in TDTs. For example, lines 25179 and 25184 had similar lifespans (93% and 95% mortality rates at 4 weeks, respectively), yet one lost significantly more fibers per TDT than the other (**Fig. 4C**).

Collectively, quantification studies strongly suggest that spontaneous fiber degeneration in flies is an aging-related process. Meanwhile, the extent of fiber degeneration in TDTs is line-specific, implying that this phenomenon is influenced by genetic factors.

## Muscle degeneration in flies is influenced by contractile activity

Extensive physical exercise has been shown to inflict damage of muscle fibers in humans [41]. To test if the level of physical activity (or mechanical stress) affects aging-related muscle degeneration in flies, we analyzed bang-sensitive flies that demonstrate abnormal neurophysiology [42]. The bang-sensitive phenotype was achieved by driving tissue-specific RNAi knockdown of *julius seizure* (*jus*) in neurons via *elav-Gal4*, as described before [32].

The experimental, bang-sensitive flies (*jus* KD^NEU^) had functional TDTs, as evident from the jump test results (**Fig. 5A**), but were prone to seizures and paralysis after a brief mechanical stimulation (*i*.*e*., vigorous shaking [32]). We aged bang-sensitive flies along with their genetically matched control (CNTR^NEU^) for 4 weeks, while applying mechanical stimulation daily. In the control flies such treatment *per se* did not promote fiber degeneration in the TDT (**Fig. 5B, D**). We noted small bright F-actin inclusions within TDTs of some flies, indicative of alterations in the fine muscle organization (**Fig. 5B, C**). These inclusions could be a sign of muscle injury, similarly to what was reported for post-exercised muscles in rats [43]. The incidence of such inclusions was notably elevated in old *jus* KD^NEU^ flies; no other tested genotype reached a similar level of F-actin inclusions (**Fig. 5C**). Importantly, bang-sensitive flies also demonstrated a small but significant increase in the levels of TDT fiber degeneration (**Fig. 5E**).

**Figure 5:**
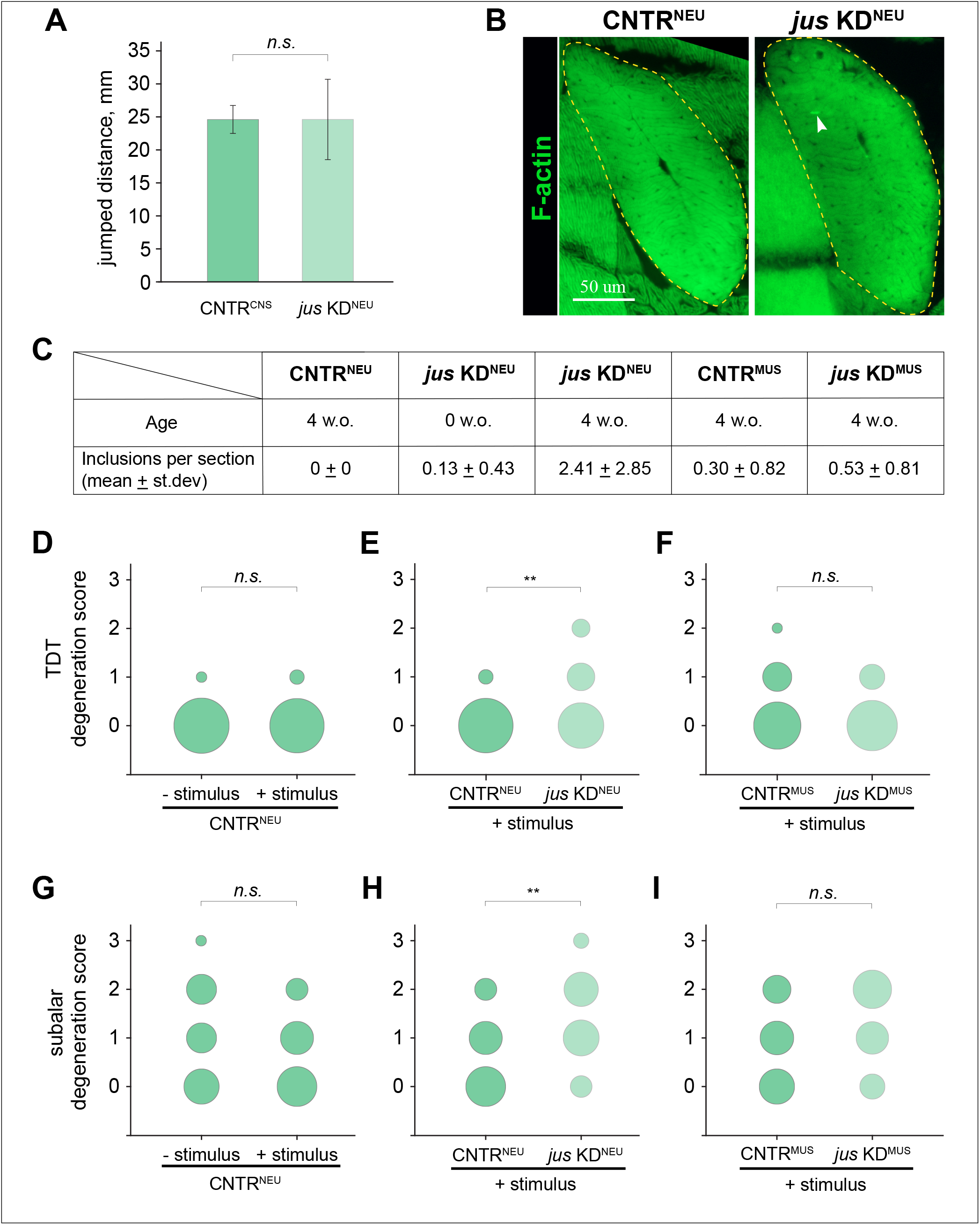
Exacerbated rates of muscle generation in bang-sensitive flies during aging. **A**. Jump test performance by flies with *jus* knockdown in neurons (*jus* KD^NEU^) and their genetically matched controls (CNTR^NEU^). 30 young flies were tested per each group. Whiskers show standard deviation. **B**. The morphology of the TDT muscle (outlined by dashed line) from control (CNTR) and experimental flies (*jus* KD^NEU^). F-actin staining; the arrowhead points at an occasional F-actin positive inclusion. Images from 4 w.o. flies are shown. **C**. Frequencies of F-actin inclusions in the TDT muscles from flies of different genotypes and ages. Each individual TDT was analyzed from two different sections and then the inclusion counts were averaged. *jus* KD^NEU^ and *jus* KD^MUS^ depict the flies with *jus* knockdown in neurons and muscles, respectively. CNTR^NEU^ and CNTR^MUS^ are their genetically matched controls. Fly age is given in weeks (w.o.) at 29°C. **D-F:** Quantification of TDT degeneration in flies of various genotypes and treatments. The area of each circle represents the frequency of muscles received each degeneration score, as per Figure 3C. Flies were aged at 29°C for 4 weeks; 30 TDT muscles were analyzed per sample. “Stimulus” indicates vigorous shaking for 10 sec daily to invoke the bang response in *jus* KD^NEU^ flies. **E-I:** Quantification of fiber degeneration in subalar muscles from the same flies and under the same conditions as described in D-F. Values of the Student’s *t-*test: * *p<0*.*05*; *** *p<0*.*001*; *n*.*s*.: *p>0*.*05*.

For additional control, we induced *jus* knockdown in muscles, using muscle-specific driver *Mef2-Gal4*, on the premise that *jus* is predominantly expressed in the nervous system but not in muscles [32]. Such flies (*jus* KD^MUS^) did not demonstrate bang sensitivity and their TDT degeneration rates were comparable with those of genetically matched control (*CNTR*^*MUS*^) (**Fig. 5C, F**).

We extended our analysis into the subalar muscle – a small, multifiber muscle immediately adjacent to the TDT (muscle 57 by *Miller* [44]). The subalar muscle belongs to the same tubular muscle type as the TDT but has a stronger SDH activity (**Fig. 2 A, D**). In general, subalar muscles demonstrated a higher level of degeneration than TDTs, but mechanical stimulation could not further exacerbate it in control, CNTR^NEU^ flies (**Fig. 5G**). Similarly to the observations made for TDTs, in bang-sensitive *jus* KD^NEU^ flies subalar muscle degeneration was more severe than in control flies **(Fig. 5H)**. Meanwhile, in bang-insensitive *jus* KD^MUS^ flies fiber degeneration in subalar muscles was not different from the control (**Fig. 5I**).

Taken together, our data demonstrate that aging-related muscle degeneration in flies is affected by the intensity of muscle activity. Our results also imply that the functional state of the nervous system is a contributing factor in muscle aging.

## Functional inactivation does not prevent muscle degeneration

To further test the contribution of physical activity to muscle degeneration in flies, we genetically decoupled the TDT muscle from neurogenic stimulation. We did that by inducing RNAi knockdown of *Ca-α1D*, which codes a subunit of the voltage-gated Ca2+ channel that is critical for action potential propagation and subsequent muscle contraction [45]. We employed the *Act79B-Gal4* genetic driver [33] to limit the effect of knockdown by the TDT muscle and thus avoid systemic paralysis and lethality in adult flies.

In the experimental knockdown flies (*Ca-α1D* KD^TDT^), TDTs developed normally and achieved the usual size and shape (**Fig. 6A**). However, the normal distribution of myonuclei in these muscles was perturbed. Instead of 2-3 central lacunae housing nuclei, the nuclei of the *Ca-α1D* KD^TDT^ flies could be found at multiple random locations scattered throughout the fiber cross-sectional area (**Fig. 6B**). Such alteration in myonuclear positioning could be indicative of changes in the fine cellular organization of the fiber. The efficacy of *Ca-α1D* genetic knockdown in the TDT was confirmed by the jump test, with 100% of the experimental flies being unable to jump (**Fig. 6C**).

**Figure 6:**
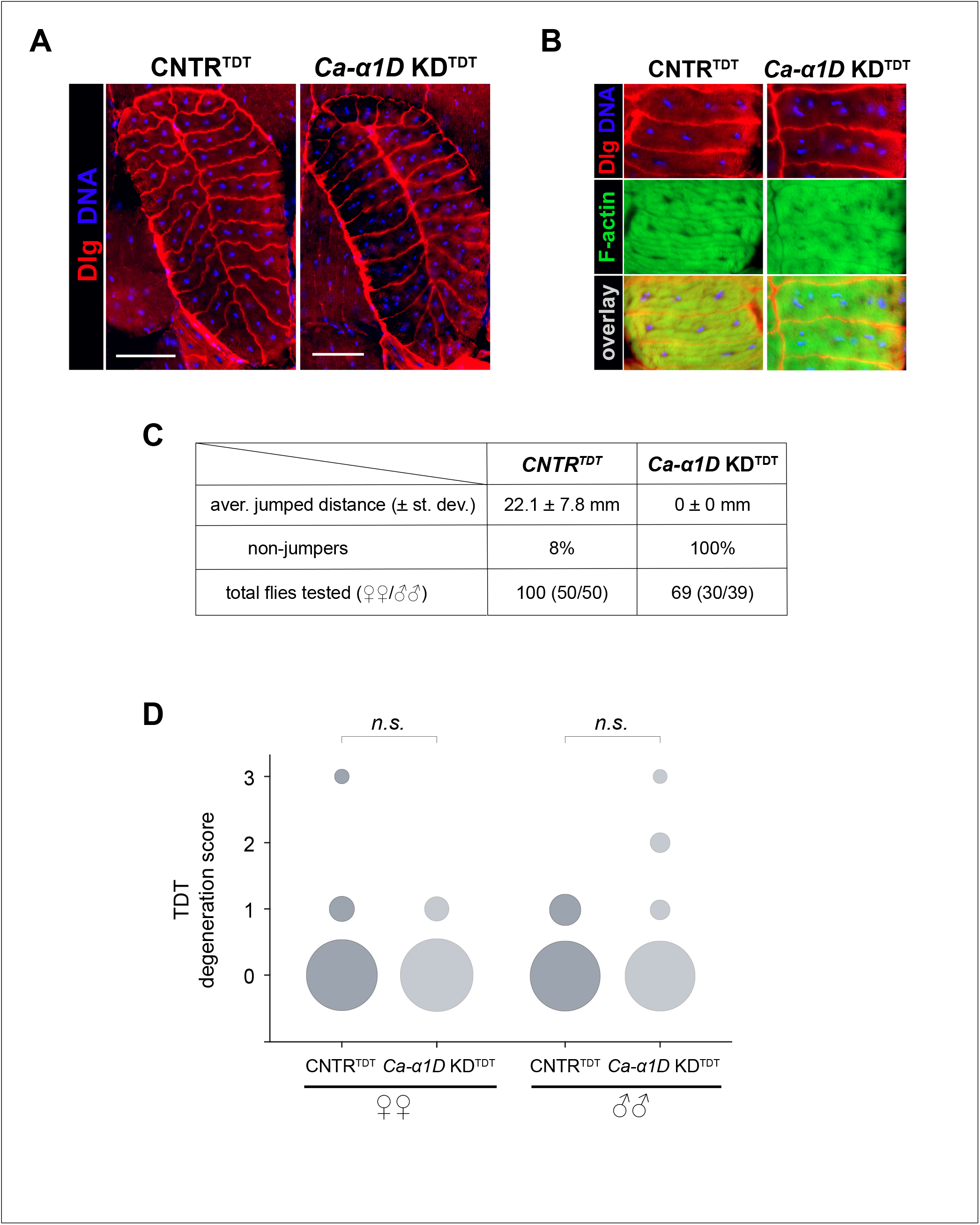
Spontaneous fiber degeneration in the TDT muscle genetically disconnected from neurogenic activation. **A**. Cross-sectional morphology of the TDT muscle. Anti-Dlg immunofluorescence (red) reveals boundaries of muscle fibers; myonuclei are counterstained with blue. No changes in the muscle size and fiber composition are evident between experimentally disabled TDT (*Ca-α1*D KD^TDT^) and a functional TDT from genetically matched control (CNTR^TDT^). **B**. Localization of myonuclei in the TDT muscle. A fragment of the TDT showing several fibers stained for F-actin (green), membranes (Dlg, red), and nuclei (blue). Distribution of myonuclei in genetically disconnected TDT (*Ca-α1*D KD^TDT^) is perturbed in comparison with control muscle (CNTR^TDT^). **C**. Validation of TDT disconnection from neurogenic stimulation via the jump test. Experimental flies (*Ca-α1*D KD^TDT^) completely lose the ability to jump. **D**. Quantification of TDT degeneration in aging flies. The area of each circle represents the fraction of muscles received a particular degeneration score, as per Figure 3C. Control (CNTR^TDT^) and experimental flies with functionally disabled TDTs (*Ca-α1*D KD^TDT^) were raised at 29°C for 4 weeks. Results for males and females are shown separately. Values of the Student’s *t-*test: *n*.*s*.: *p>0*.*05*.

Aging *Ca-α1D* KD^TDT^ flies demonstrated low frequency of TDT degeneration that was detected in both sexes and was not statistically different from the level of TDT degeneration observed in age-matching control flies (CNTR^TDT^) (**Fig. 5D**).

Since muscles with disabled contractile activity remain sensitive to spontaneous degeneration, we conclude that this phenomenon is aging-related and may depend on intrinsic muscle factors. In addition, we reveal that the fine fiber organization is dependent on the functional state of the TDT muscle.

## DISCUSSION

This study characterizes aging-related changes in the somatic musculature of adult *Drosophila*. We reveal that all somatic muscle types are prone to spontaneous degeneration in aging flies. By using quantitative analysis, we demonstrate that muscle damage correlates with aging (as determined by chronological, functional, and populational criteria) and it is influenced by genetic background and physical activity. Historically, muscle morphology in the *Drosophila* model was used primarily to assess effects of genetic mutations or factors on muscle development [24, 25, 46] or to study experimentally induced muscle degeneration [26, 27]. By comprehensively describing the normal aging-related muscle degeneration, we lay the foundation for using *Drosophila* in systematic genetic screening for identifying factors that affect the quality of muscle aging. We believe that the methodology developed in this study, coupled with the genetic proficiency of the *Drosophila* model, will accelerate discoveries of novel targets for sarcopenia research.

### The mechanism of muscle degeneration in adult flies

Instances of muscle death under normal conditions in *Drosophila* have been previously reported as “programmed muscle death”. Most larval muscles undergo rapid degeneration in response to hormonal cues during metamorphosis in pupae or shortly after it, in young adults [47–49]. Under this scenario, fiber degeneration is initiated from within via intracellular signaling that activates effector caspases (in apoptosis) or lysosomes (in autophagy) [47, 50]. Similar regulations take place in anuran larval muscles (within the tadpole tail) or in mammalian muscles upon denervation or disuse [51, 52].

There is a substantial difference between the programmed death of larval muscles and the spontaneous degeneration of adult muscles reported in this study. Doomed larval muscles lose attachment, round up, and undergo fragmentation of their sarcoplasm and nuclei [48]. Interestingly, the fragmented sarcoplasm retains positive staining for polymerized actin [47, 48]. Similar nuclear fragmentation was reported for dying fibers of denervated mammalian muscle [53]. However, none of these transitional changes characterize the degenerate adult muscles of our study, as they retain the original shape while completely losing polymerized actin. Moreover, in some cases we could observe segmental degeneration affecting only a local area of the fiber (**Fig. 2E**), which would not be possible in the programmed muscle death since it affects the entire muscle fiber. Another striking difference is that the tubular network (revealed by anti-Dlg immunostaining) is completely lost in degenerate adult fibers (**Fig. 2C**), while it lasts in dying larval muscles [47].

Taken together, our observations suggest that adult muscles mainly die not because of invoked apoptotic or autophagy program, but rather via necrosis. Necrosis is a form of cell death that is caused by a loss of cellular integrity due to the damage of plasma membrane and does not depend on an intracellular regulation. Consequently, fiber necrosis, especially segmental necrosis, occurs in skeletal musculature of humans and laboratory animals after extensive exercise [43, 54, 55]. Necrotic fibers are also commonly present in human patients with muscular dystrophies and some myopathies [56–58]. Usually, the presence of necrotic fibers attracts phagocytes [43, 58]. As will be noted below, we did not observe accumulation of cellular masses at the sites containing degenerating fibers. This apparent lack of cellular immune response to the presence of dead fibers could be explained by a diminished reactivity of the immune system in adult flies [59].

### No damage repair in Drosophila muscles, bad or good?

Adult somatic muscles in *Drosophila* are traditionally viewed as lacking structural plasticity and regenerative capacity. The latter was recently challenged with a study describing novel adult muscle stem cells in flies [60]. However, during our study, we could not find signs of muscle repair. Regeneration signatures of injured mammalian muscle include masses of mononucleated cells concentrating around the sites of damage [61]. Indeed, even a small segment of muscle fiber (*e*.*g*., TDT) contains tens to hundreds of nuclei and would require an equal number of mononucleated progenitors for repair. However, we did not detect swarms of nuclei around degenerating muscle fibers. We also did not observe intact nuclei *within* degenerating fiber segments, which excludes a putative repair of muscle fibers by endoreplication mechanisms as seen in regenerating cardiac muscle [62]. Therefore, our data support the canonical view, according to which regeneration in adult *Drosophila* muscles is rare. Although this fact separates fly muscles from regeneration-potent vertebrate muscles, it gives a pragmatic advantage to our model: the lack of regenerative capacity enables a lasting record of muscle damage, making the identification of factors affecting structural integrity of muscles more straightforward.

### The effect of mechanical force on muscle architecture

Disconnected, dysfunctional muscle fibers undergo subtle but noticeable changes in their morphological appearance. Denervated muscle fibers in mammals and humans have reduced sizes and assume atypical, angular shape on the cross section; upon long-term denervation, fibers with centrally located myonuclei arise [63]. In *Drosophila*, physical connection to the nervous system is important in early steps of myogenesis as it can determine the type and final size of the developing muscle [64]. However, there is much less known about the role of neuronal connection in the maintenance of fully formed adult *Drosophila* muscles, considering the low structural plasticity of the latter [33]. In this study, we demonstrate that functional disconnection of the TDT muscle from neurogenic stimulation affects the nuclear positioning and results in a scattered pattern of myonuclei.

In mammals, myonuclei move between peripheral and central regions of the muscle fiber during development or regeneration [65]. Nuclear movement is important for normal muscle functioning since its misregulation leads to centronuclear myopathies in humans [66, 67]. *Drosophila* was instrumental in dissecting the genetic control of myonuclear positioning, revealing the key factors participating in this process [68, 69].

However, the gene *Ca-α1D* that was targeted by RNAi in our experiments has not been previously associated with myonuclear control. It is possible, therefore, that the nuclear misalignment within dysfunctional TDTs is a result of non-genetic factors. Indeed, mechanical forces have been shown to determine myonuclear positioning within mammalian muscle fibers [70]. It is intriguing to hypothesize that contractile forces determine the nuclear localization within the TDT fibers as well, despite some architectural differences in the organization of *Drosophila* [33] and mammalian [71] muscle fibers.

### The role of the nervous system in muscle aging

Our observations imply that mechanical damage could be a major driver of muscle degeneration in aging flies. The nervous system regulates the intensity and duration of muscle contractions and may modulate the amount of mechanical stress received by muscles. Using seizure-prone, bang-sensitive flies, we demonstrated how a compromised nervous system can affect and promote muscle degeneration. A coincidental muscle damage from seizures was also reported in humans [72]. Although seizures are the extreme stimulus to inflict muscle damage, we hypothesize that even subtle deviations from the normal neurogenic activation of muscles would influence wear-and-tear of muscle fibers, if continuously occurring over the lifetime.

How relevant is this finding in the context of mammalian muscle? Upon acute injury, mammalian muscle can effectively repair damaged fibers and replace dead fibers [13]. However, chronic or recurring damage can eventually overwhelm the regenerative capacity, as seen in the case of Duchenne’s and other progressive muscle dystrophies [15]. Furthermore, the efficacy of muscle regeneration may decline with age [14], potentially leading to the reduction of fiber counts reported for older adults [11].

Traditionally, genetic factors affecting functioning of the nervous system are not considered *bona fide* candidates for sarcopenia [18], although neurogenic etiology of sarcopenia has been proposed [73, 74]. Embracing the potential contribution of the nervous system to muscle degeneration will substantially expand the range of candidates for sarcopenia research.

## ACKNOWLEDGEMENTS

This work was supported by grant R03 AG048496, awarded to ALB by the NIH. The authors are grateful for the flies provided by Dr. Aaron Johnston. The integrin monoclonal antibodies, developed by Dr. Brower, were obtained from the Developmental Studies Hybridoma Bank (dshb.biology.uiowa.edu), created by the NICHD of the NIH and maintained at the University of Iowa. The authors would like to acknowledge Kennesaw State University Academic Affairs and Georgia BIO for support of the Microscopy Core Facility, where most of imaging was performed.

## Notes

### Competing Interest Statement

The authors have declared no competing interest.

